# IMPDH polymers accommodate both catalytically active and inactive conformations

**DOI:** 10.1101/152173

**Authors:** Sajitha Anthony, Anika L. Burrell, Matthew C. Johnson, Krisna C. Duong-Ly, Yin-Ming Kuo, Peter Michener, Andrew Andrews, Justin M. Kollman, Jeffrey R. Peterson

## Abstract

Several metabolic enzymes undergo reversible polymerization into macromolecular assemblies. The function of these assemblies is often unclear but in some cases they regulate enzyme activity and metabolic homeostasis. The guanine nucleotide biosynthetic enzyme inosine monophosphate dehydrogenase (IMPDH) forms octamers that polymerize into helical chains. In mammalian cells, IMPDH filaments can associate into micron-length assemblies. Polymerization and enzyme activity are regulated in part by binding of purine nucleotides to an allosteric regulatory domain. ATP promotes octamer polymerization, whereas GTP promotes a compact, inactive conformation whose ability to polymerize is unknown. An open question is whether polymerization directly alters IMPDH catalytic activity. To address this, we identified point mutants of human IMPDH2 that either prevent or promote polymerization. Unexpectedly, we found that polymerized and non-assembled forms of IMPDH have comparable catalytic activity, substrate affinity, and GTP sensitivity and validated this finding in cells. Electron microscopy revealed that substrates and allosteric nucleotides shift the equilibrium between active and inactive conformations in both the octamer and the filament. Unlike other metabolic filaments, which selectively stabilize active or inactive conformations, IMPDH filaments accommodate multiple states. Thus, although polymerization alone does not impact catalytic activity, substrate availability and purine balance dramatically affect IMPDH filament architecture.

## INTRODUCTION

An increasing number of non-cytoskeletal proteins undergo reversible polymerization in cells and in vitro. Indeed, large scale image-based or proteomic screens have identified hundreds of proteins that dynamically form assemblies in response to nutrient stress (Alberti *et al.*, 2009; Narayanaswamy *et al.*, 2009; Noree *et al.*, 2010; O'Connell *et al.*, 2012; O'Connell *et al.*, 2014; Aughey and Liu, 2015; Liu, 2016; Shen *et al.*, 2016). Interestingly, this phenotype is disproportionately associated with metabolic enzymes, although the reason for this is unclear (O'Connell *et al.*, 2012). In many cases the functional role of polymerization has not yet been determined, but for the few metabolic filaments that have been characterized, polymerization regulates enzyme activity and is important for maintaining homeostasis (Beaty and Lane, 1983; Barry *et al.*, 2014; Petrovska *et al.*, 2014; Lynch *et al.*, 2017).

Inosine monophosphate dehydrogenase (IMPDH) reversibly assembles into helical polymers of stacked octamers in an ATP-dependent manner in vitro (Labesse *et al.*, 2013) and into larger filamentous bundles in cells (Ji *et al.*, 2006). This enzyme catalyzes the rate-limiting step in *de novo* guanine nucleotide biosynthesis, nicotinamide adenine dinucleotide (NAD^+^)-dependent oxidation of IMP into xanthine monophosphate (Hedstrom, 2009). Humans possess two differentially expressed IMPDH genes (IMPDH1 and 2) sharing 84% amino acid sequence identity (Carr *et al.*, 1993) and both can assemble into filaments (Gunter *et al.*, 2008; Thomas *et al.*, 2012). Although substrate channeling has been implicated in nucleotide biosynthesis, IMPDH filaments have not been shown to colocalize with other purine biosynthesis enzymes. However, the pyrimidine biosynthetic enzyme CTP synthase (CTPS) co-assembles with IMPDH in some circumstances (Keppeke *et al.*, 2015). This suggests the potential for coordination of purine and pyrimidine biosynthetic pathways although the role of IMPDH polymerization in this regulation has remained obscure. Filaments of bacterial CTPS are inhibited while eukaryotic filaments are active, in both cases due to stabilization of specific conformational states in the polymer, and this has been proposed as a general mechanism by which metabolic filaments may regulate enzyme activity (Barry *et al.*, 2014; Strochlic *et al.*, 2014; Lynch *et al.*, 2017).

A major challenge in assessing the impact of assembly on IMPDH catalytic activity in cells is the fact that IMPDH is generally not polymerized in cultured cell lines grown in rich media. Filament assembly can be triggered, however, by treating cells with inhibitors of purine biosynthesis, including the IMPDH inhibitors mycophenolic acid or ribavirin, inhibitors of enzymes acting either upstream or downstream of IMPDH (e.g. 6-diazo-5-oxo-L-norleucine and decoyinine) (Carcamo *et al.*, 2014), or by depleting the culture media of essential purine precursors such as glutamine or inhibiting one carbon metabolism (Calise *et al.*, 2014; Calise *et al.*, 2016). This responsiveness of IMPDH to reduced flux through the guanine biosynthetic pathway suggests that assembly could be associated with a homeostatic mechanism to restore guanine nucleotide levels. Alternatively, IMPDH filaments could serve as a storage depot for inactive enzyme. These experimental conditions, however, dramatically affect IMPDH activity independently of filament assembly, confounding their use to assess filament catalytic activity. We, therefore, sought to develop approaches to address the effect of IMPDH polymerization on catalytic activity definitively. Here, using novel IMPDH2 point mutants, electron microscopy, biochemical assays and isotopic tracer studies in live cells, we demonstrate that IMPDH polymerization alone does not alter its catalytic activity, and that allosteric regulation of enzyme activity occurs in both the free enzyme and the polymer.

## RESULTS AND DISCUSSION

### Identification of polymerization altering mutants of human IMPDH2

We sought to identify residues at the putative octamer polymerization interface that when mutated to alanine would disrupt IMPDH2 polymerization. The crystal structure of human IMPDH2 (PDB code:1NF7) contains stacked octamers that resemble IMPDH filaments (Labesse *et al.*, 2013). On the assumption that these crystal contacts are the same as polymerization interfaces in IMPDH filaments, we designed a series of mutants that we predicted would block polymerization (Figure 1A). Individual myc-tagged alanine mutants were transiently transfected in HEK293 cells (Figure 1B) and filament assembly was induced by overnight treatment with the IMPDH inhibitor mycophenolic acid (MPA) and visualized by immunostaining with anti-myc antibodies. Among other amino acids examined (threonine 10 and aspartate 15), only alanine substitution of tyrosine 12 (Y12A) or arginine 356 (R356A) potently disrupted filament induction by MPA while wild type assembled as expected (Figure 1C). These residues are conserved across vertebrate IMPDH isoforms, supporting their functional importance.

**Figure 1:**
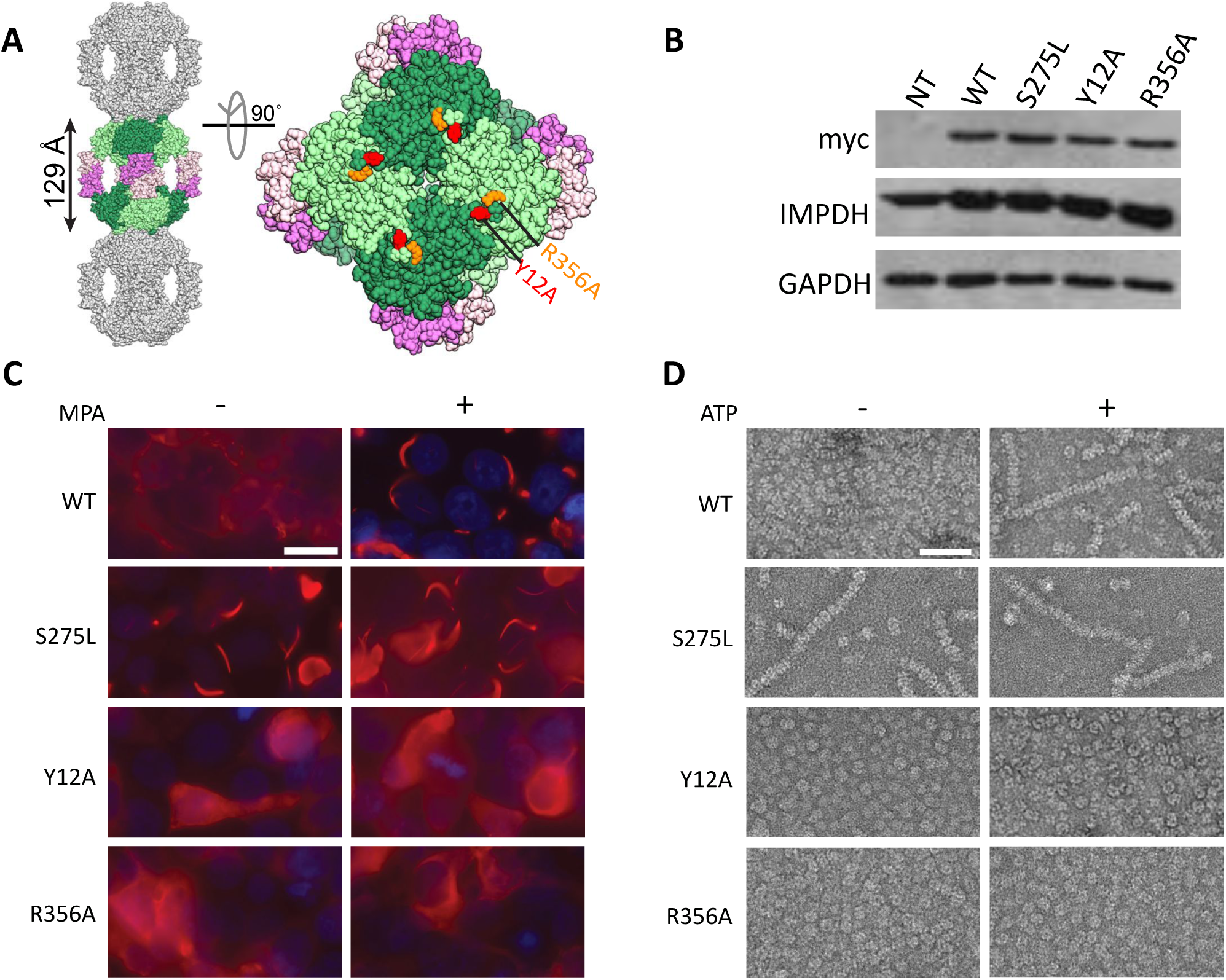
Identification of filament disrupting and promoting point mutations of human IMPDH2. (A) The human IMPDH2 octamer (PDB file 1NF7). One octamer (color: greens - catalytic domains, pinks - regulatory Bateman domains) is shown with a subset of crystal packing neighbors (gray). Residues tyrosine 12 (red) and arginine 356 (orange) located at crystal contact sites near the presumed filament assembly interface are indicated. (B) Immunoblot of HEK293 cell lysates with antibodies directed against myc-tagged IMPDH2, total IMPDH, or loading control glyceraldehyde 3-phosphate dehydrogenase (GAPDH). (C) Immunofluorescence images of HEK293 cells transiently transfected with the indicated IMPDH2-myc construct stained with anti-myc antibodies. Cells were either treated with 10 μM mycophenolic acid to induce IMPDH2 assembly or not, as indicated. Images are representative of 3 experiments. Scale bar, 20 μm. (D) Negative stain electron microscopy of purified, recombinant IMPDH2 proteins incubated in the presence of 5 mM NAD+, with and without 1 mM ATP. Images represent 2 separate experiments. Scale bar, 50 nm.

To confirm that these mutations were acting directly on IMPDH2 self-assembly (as opposed to disrupting protein folding or interactions with assembly partners), we used electron microscopy to directly visualize recombinant wild type and mutant IMPDH2 (Supplemental Figure 1). The wild type protein appears as a mixture of octamers and tetramers with occasional short filaments in the absence of ligands, and formed long polymers in the presence of ATP (Figure 1D). In the absence of ligands, Y12A and S356A appear as a mixture of octamers and tetramers without any short polymers, and no polymers were observed in the presence of ATP, indicating that the mutations directly interfere with polymerization (Figure 1D).

A study of human IMPDH sequence polymorphisms in healthy individuals identified a rare, non-synonymous single nucleotide polymorphism altering serine 275 to leucine (S275L) in IMPDH1 (Wu *et al.*, 2010) that constitutively assembled into filaments when expressed in cells (Wu, 2011). We engineered this mutation into IMPDH2 and confirmed the propensity of S275L to assemble in transfected cells, even in the absence of MPA (Figure 1C). Unlike wildtype protein, purified recombinant S275L robustly assembles polymers in both the presence and absence of ATP, indicating that the propensity of S275L to polymerize is an intrinsic consequence of the mutation (Figure 1D). S275L is localized adjacent to the NAD^+^ binding site and the basis for its constitutive filamentation is unclear. Nevertheless, this construct served as a useful experimental counterpoint to the non-assembling mutants.

### Catalytic activity of IMPDH2 mutants

Next, we assessed the catalytic activity of purified, recombinant wild type and mutant IMPDH2 by monitoring NADH production in real time by fluorescence (Figure 2A & 2B). Initial reaction rates were measured for unpolymerized IMPDH under conditions of saturating substrates NAD^+^ and IMP in the absence of ATP. We observed no significant difference in the specific activity of wild type and any of the three IMPDH2 mutants (Figure 2C). We next measured the activity of each variant in the absence or presence of ATP, which promotes polymerization of wild type IMPDH2 but not Y12A and R356A (Figure 1D). We observed no evidence for increased activity of wild type IMPDH2 under polymerization conditions, consistent with reports that ATP does not enhance IMPDH activity (Mortimer and Hedstrom, 2005; Pimkin and Markham, 2008; Labesse *et al.*, 2013; Buey *et al.*, 2015) (Figure 2D). Similarly, non-polymerizing IMPDH2 mutants were unaffected by ATP. Taken together, the comparable activity of IMPDH2 under polymerizing and non-polymerizing conditions indicates that assembly of purified IMPDH2 does not affect specific activity.

**Figure 2:**
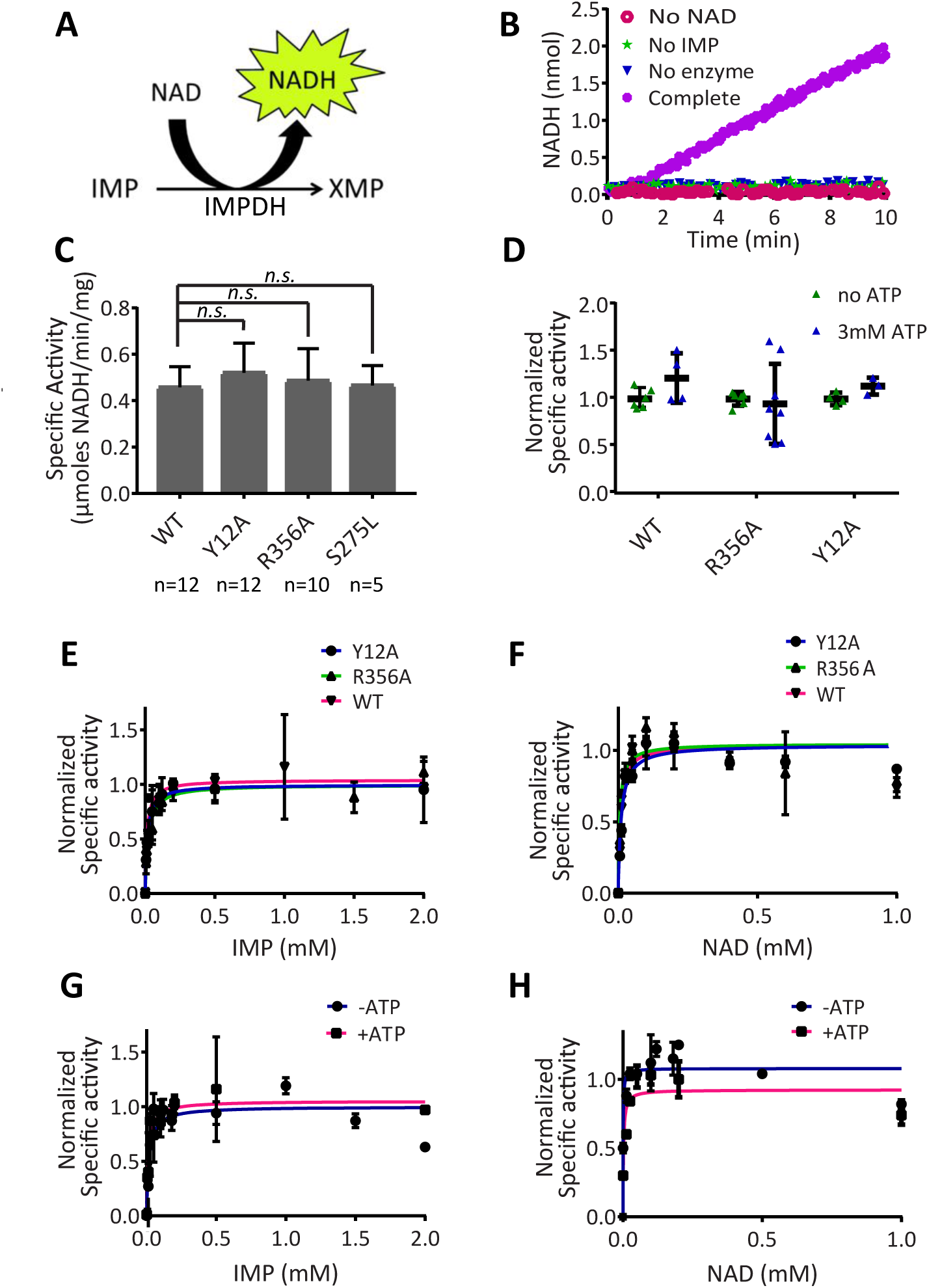
IMPDH2 polymerization does not alter its catalytic activity in vitro. (A) Schematic of the IMPDH reaction illustrating production of NADH, which can be detected by fluorescence. (B) NADH fluorescence was monitored in reactions lacking NAD+, IMP, IMPDH (no enzyme), or in a reaction containing all reaction components (Complete). (C) IMPDH activity was measured for wild type and mutant forms under non-assembling conditions. All mutants showed no significant difference in activity compared to wild type (two-sided Student’s t test; P > 0.05). Bars denote mean and standard deviation. (D) IMPDH activity assays as in (C) were conducted in the absence or presence of ATP to induce assembly. ATP did not significantly alter activity in any case (two-sided Student’s t test). (E & F) Initial reaction rates are plotted as a function of substrate. concentration for IMP or NAD+ comparing wild type IMPDH2 with non-assembling mutants under polymerization conditions (1 mM ATP) or for wild type in the presence or absence of ATP (G & H). Michael is constants determined from these titrations are given in Supplemental Table 1. 2-5 replicates per substrate concentration.

Measuring catalytic activity under saturating substrate concentrations might mask subtle differences in IMPDH2 substrate affinity in the polymer compared with free enzyme. We therefore determined apparent Michaelis constants (K_m_) of mutant and wild type IMPDH2 for substrates IMP and NAD^+^ by measuring initial reaction velocity at a range of substrate concentrations under polymerizing and non-polymerizing conditions. Mutant and wild type IMPDH2 affinity for IMP (Figure 2E) and NAD^+^ (Figure 2F) were indistinguishable (see also Supplemental Table 1), implying that wild type polymers and unassembled mutants have comparable substrate affinities. Consistent with this, we found that triggering wild type IMPDH2 polymerization with ATP also did not affect IMP affinity (Figures 2G). A 10-fold increase in the Km(app.) for NAD^+^ is likely due to competitive binding of ATP to the NAD^+^ site rather than an effect of polymerization since no difference in NAD^+^ substrate affinity was observed comparing wild type and non-polymerizing mutants (Figure 2F). We conclude that substrate affinity is not substantially altered by IMPDH2 polymerization. More broadly, these results demonstrate that Y12A and R356A are true separation-of-function mutations, abolishing the polymerization ability of IMPDH2 while maintaining its catalytic activity.

### IMPDH assembly does not alter guanine nucleotide biosynthetic flux in vivo

Next, we considered whether IMPDH2 polymerization might alter its catalytic activity in the cellular context in a way not detectable for the purified protein in vitro, for example by regulating IMPDH2 interaction with other proteins. We therefore established a method to monitor IMPDH activity in live cells by monitoring incorporation of isotopically labeled glycine, provided in the culture media, into GMP, a downstream metabolic product of IMPDH, or AMP as a control. Because IMPDH is rate-limiting for guanine nucleotide synthesis, we reasoned that alterations in IMPDH activity would impact labeled glycine incorporation into the GMP pool.

We focused our analysis on comparing the activity of the constitutively polymerizing S275L with the non-polymerizing Y12A. Importantly, these forms exhibit identical catalytic parameters *in vitro* (Figures 2, C-H). HEK293 cells were transfected with each construct or empty vector and were immunostained with antibodies against IMPDH or anti-myc antibodies specific to the transfected proteins. Total IMPDH expression was increased by ~ 2-fold in IMPDH2-transfected cells compared to empty vector-transfected cells, evident from the increased immunofluorescence signal (Figure 3A) and by immunoblot (Figure 3B). S275L and Y12A expressed at similar levels. As expected, S275L-transfected cells showed robust IMPDH2 filaments whereas Y12A-expressing cells showed diffuse, cytoplasmic staining (Figure 5A).

**Figure 3:**
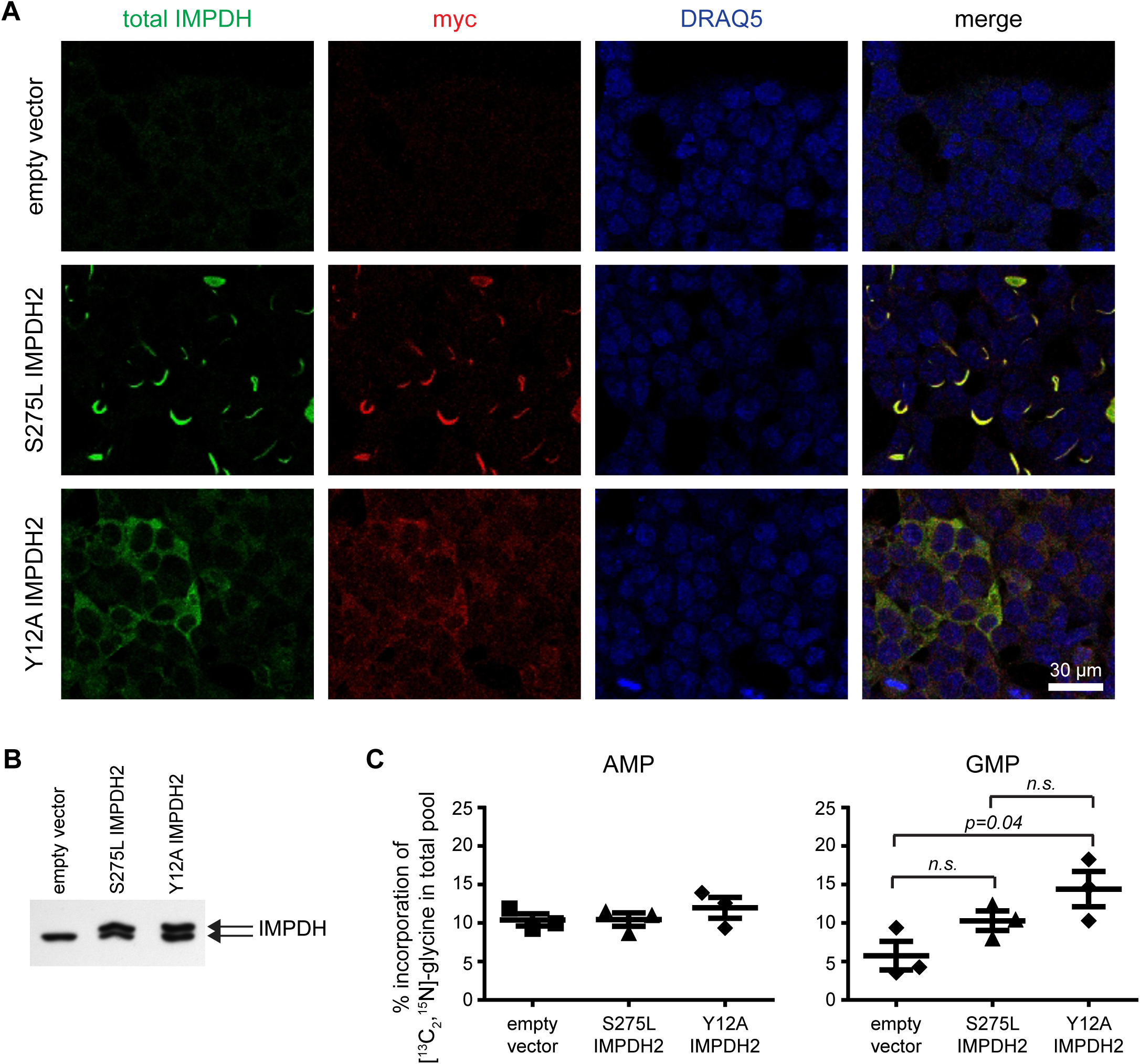
IMPDH filament assembly does not alter guanine nucleotide biosynthesis. (A) HEK293 cells transfected with either empty vector, S275L IMPDH2-myc or Y12A IMPDH2-myc were visualized with antibodies against myc, total IMPDH, or a DNA stain (DRAQ5). S275L IMPDH2 assembles into filaments while Y12A remains diffusely cytoplasmic. (B) Representative immunoblot of total IMPDH expression in transfected HEK293 cells. The higher molecular weight band is the transfected protein whereas the lower band is endogenous IMPDH. (C) Incorporation of [^13^C_2_, ^15^N]-glycine into AMP and GMP. Bars denote mean and standard error for three biological replicates conducted on different days. No statistical differences (two-sided Student’s t test; P > 0.05) were found between samples for incorporation of [13C2, 15N]-glycine into AMP pools.

Parallel cultures of transfected cells were incubated with [^13^C2, ^15^N]-glycine at a 3-fold molar excess over unlabeled glycine in the medium. This resulted in a three unit mass increase in glycine-derived metabolites, allowing a robust quantitation of pulse-labeled AMP/GMP by ultra performance liquid chromatography-tandem mass spectrometry (UPLC-MS/MS). Unlike the percentage of isotopically labeled AMP, the isotopically labeled GMP fraction was modestly increased in both IMPDH2-transfected cells compared to empty vector control (reaching statistical significance only for Y12A), consistent with an increase in total IMPDH activity (Figure 3C). Importantly, however, the percent of GMP incorporating labeled glycine, and therefore synthesized via IMPDH, was indistinguishable in S275L- and Y12A-expressing cells, demonstrating comparable flux through IMPDH regardless of its polymerization state. This finding validates our in vitro results and shows that diffuse and filamentous forms of IMPDH2 share comparable catalytic activity in vivo.

The unexpected similarity in the activity of diffuse and filamentous IMPDH2 begs the question of why IMPDH assembles in cells under conditions of limited nucleotide biosynthetic flux. Our methods may not detect subtle differences in the kinetics of transitions between the expanded and compressed states mediated by cooperative transitions within the polymer that could be relevant in vivo. Alternatively, filament assembly could fulfill a different function such as sequestering IMPDH from non-catalytic functions (Kozhevnikova *et al.*, 2012), or scaffolding interactions with other proteins. The present work introduces separation-of-function mutants of IMPDH that will be useful for addressing the role of assembly in vivo.

### GTP effects on IMPDH octamer conformation and catalytic activity

Conformational changes in the IMPDH octamer have been associated with altered catalytic activity. For example, a recent study found that eukaryotic IMPDHs are subject to feedback inhibition by low millimolar GDP or GTP (Buey *et al.*, 2015). The crystal structure of *Ashbya gossypii* IMPDH bound to GDP (PDB: 4Z87) revealed a collapsed conformation of the IMPDH octamer mediated by the binding of three GDP molecules to sites within the Bateman domain (Buey *et al.*, 2015). Two sites overlap with the canonical Bateman domain ATP-binding sites, implying that competition between adenine and guanine nucleotides for these allosteric regulatory sites and/or binding of a guanine nucleotide to the third site can trigger a dramatic conformational change in the IMPDH octamer from an expanded, catalytically active form to a collapsed, inactive form. Residues that mediate GTP binding are conserved in eukaryotic but not prokaryotic IMPDHs, though the ability of human IMPDH to adopt the collapsed state has not been reported. To examine this directly we carried out a comprehensive investigation of IMPDH2 conformational states using negative stain and cryo-electron microscopy. We sought to determine which conformations the human octamer can adopt in the free and filament contexts and what role ligands play in determining these conformations.

We explored a range of combinations and concentrations of ATP, GTP, and substrates and assessed the structural consequences using negative stain images and two-dimensional, reference-free classification of IMDPH2 filaments. We identified four major classes of filament segments: expanded, collapsed, bent, and ‘poorly aligning’ (Figure 4, A-D). The spacing between octamer centers in the expanded conformation is ~110 Å compared to ~95 Å in the collapsed conformation. The bent conformation is well-resolved with one side matching the spacing of the collapsed conformation and the other side matching the expanded conformation, suggesting that in some cases octamers may occupy conformational states intermediate in the transition between expanded and collapsed, with IMPDH2 monomers present in mixed states within a single octamer. We defined as 'poorly aligning' segments that were assigned to classes without detailed features. This finding demonstrates the surprising plasticity of the filaments to accommodate both expanded and collapsed octamer conformations. We quantified the frequency of these classes in the presence or absence of ATP, GTP and substrates (Figure 4B). This analysis shows that binding of either substrate shifts the conformational equilibrium toward the expanded octamer, and that GTP promotes the collapsed conformation, suggesting that the balance of GTP and substrates tunes the conformational state of the IMPDH filament.

**Figure 4:**
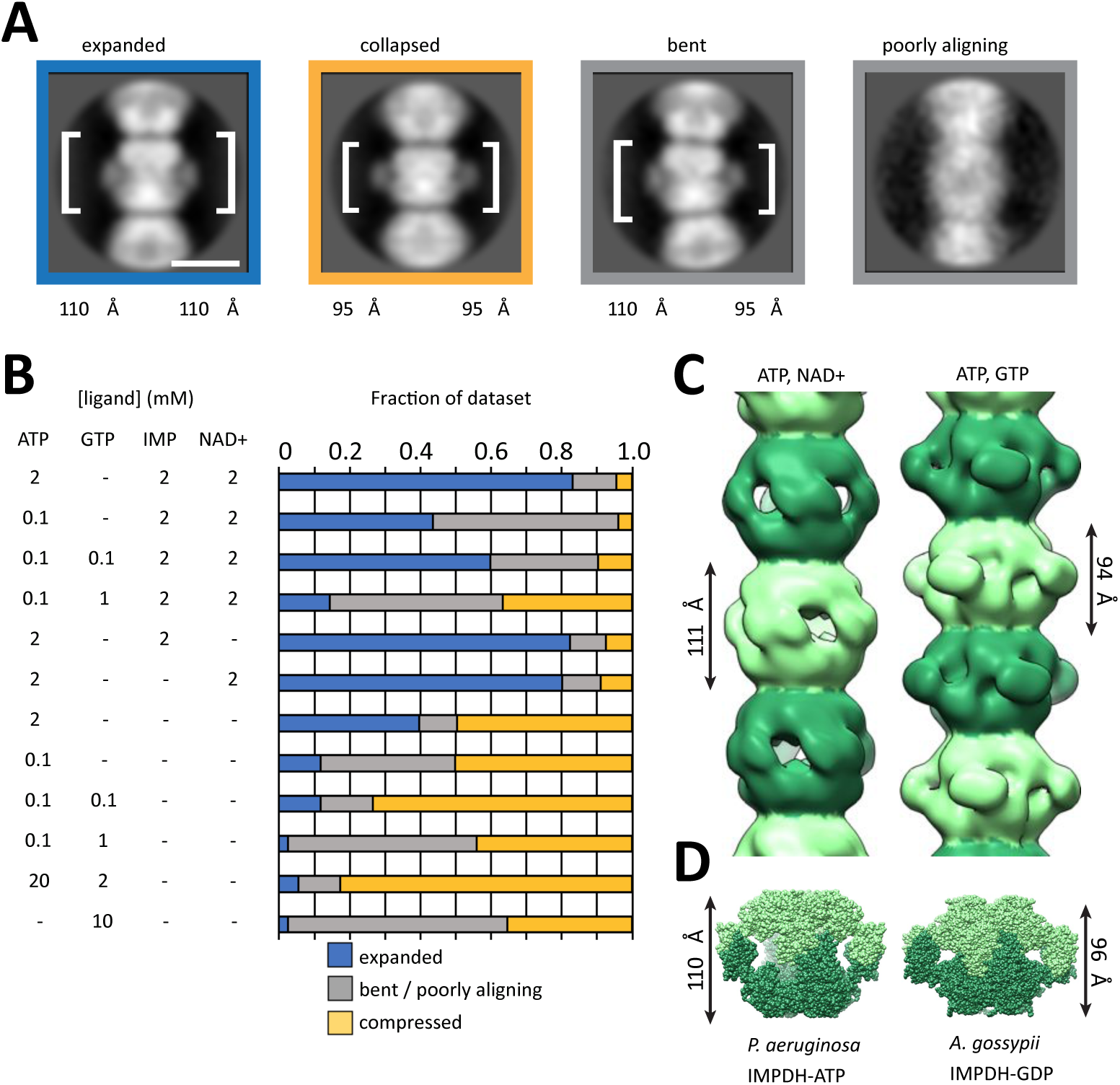
Ligands influence IMPDH filament architecture by altering the conformation of IMPDH protomers. (A) Representative class averages of IMPDH filaments in four different conformational states, with the height of one IMPDH octamer indicated. (B) Quantification of the fraction of IMPDH helical segments in each conformational state as a function of ligand concentrations. (C) Negative stain EM reconstruction of IMPDH filament structures in the expanded substrate-bound (0.1 mM ATP, 2 mM NAD+) and collapsed GTP-bound (0.1 mM ATP, 0.1 mM GTP) conformations. The refined helical rotation are rise for the expanded filament are 30° and 111 Å, and for the collapsed filament 35.5° and 94 Å, consistent with the rise measured directly from two-dimensional class averages in (A). (D) The heights of crystal structures of *P. aeruginosa* IMPDHATP and *A. gossypii* IMPDH-GDP closely match the refined helical rise of the human IMPDH filaments in the expanded and collapsed states, respectively.

We performed three-dimensional reconstructions of IMPDH2 filaments under conditions where the equilibrium was shifted most strongly toward the expanded conformation (0.1 mM ATP, 3 mM IMP, 5 mM NAD^+^) or the collapsed conformation (0.1 mM ATP, 0.1 mM GTP) (Figure 4C), yielding structures at ~20 Å resolution. The structures reveal helical polymers formed from stacked octamers that differ significantly from each other in their refined helical rise (111 Å for expanded and 95 Å for the collapsed state), consistent with the 2-dimensional class averages. In the collapsed conformation there is a 30.5° rotation between octamers and the collapsed filament exhibits an additional 5° of rotation between octamers. It appears that the filament assembly contacts between octamers are relatively fixed, and the differences in helical symmetry arise primarily from differences in the connection between the regulatory Bateman domain and the catalytic domain.

Strikingly, neither filament conformation matches the ~130 Å spacing of the human IMPDH2 octamer in its crystal lattice (Figure 1A). Rather, the expanded conformation most closely resembles the ATP-bound active octamer conformation first reported for *Pseudomonas* IMPDH (Labesse *et al.*, 2013), while the collapsed conformation is most similar to the *Ashbya* GDP-bound inhibited form (Buey *et al.*, 2015) (Figure 4D). The identification of conditions that shift the conformation of IMPDH between collapsed and expanded conformations suggests the potential for cooperativity in the transition of polymerized octamers between states. This model would imply that polymers may function to coordinate octamer conformational changes *en masse*.

We sought to confirm that GTP-bound human IMPDH2 adopts a collapsed conformation similar to *Ashbya* IMPDH with high-resolution cryo-electron microscopy. We determined the 8.7 Å structure of GTP-bound Y12A IMPDH2 (Figure 5A). At this resolution alpha-helices are well-defined, allowing us to model the overall conformation of the protein by rigid-body fitting of the human IMPDH catalytic and Bateman domains into the EM density (Figures 5B & 5C). Comparison of this atomic model to the *Ashyba* GDP-bound IMPDH2 crystal structure (PDB ID 4Z87) unambiguously reveals that under these conditions Y12A assumes the collapsed conformation (Figure 5D) and not the extended form (Figure 5E).

**Figure 5:**
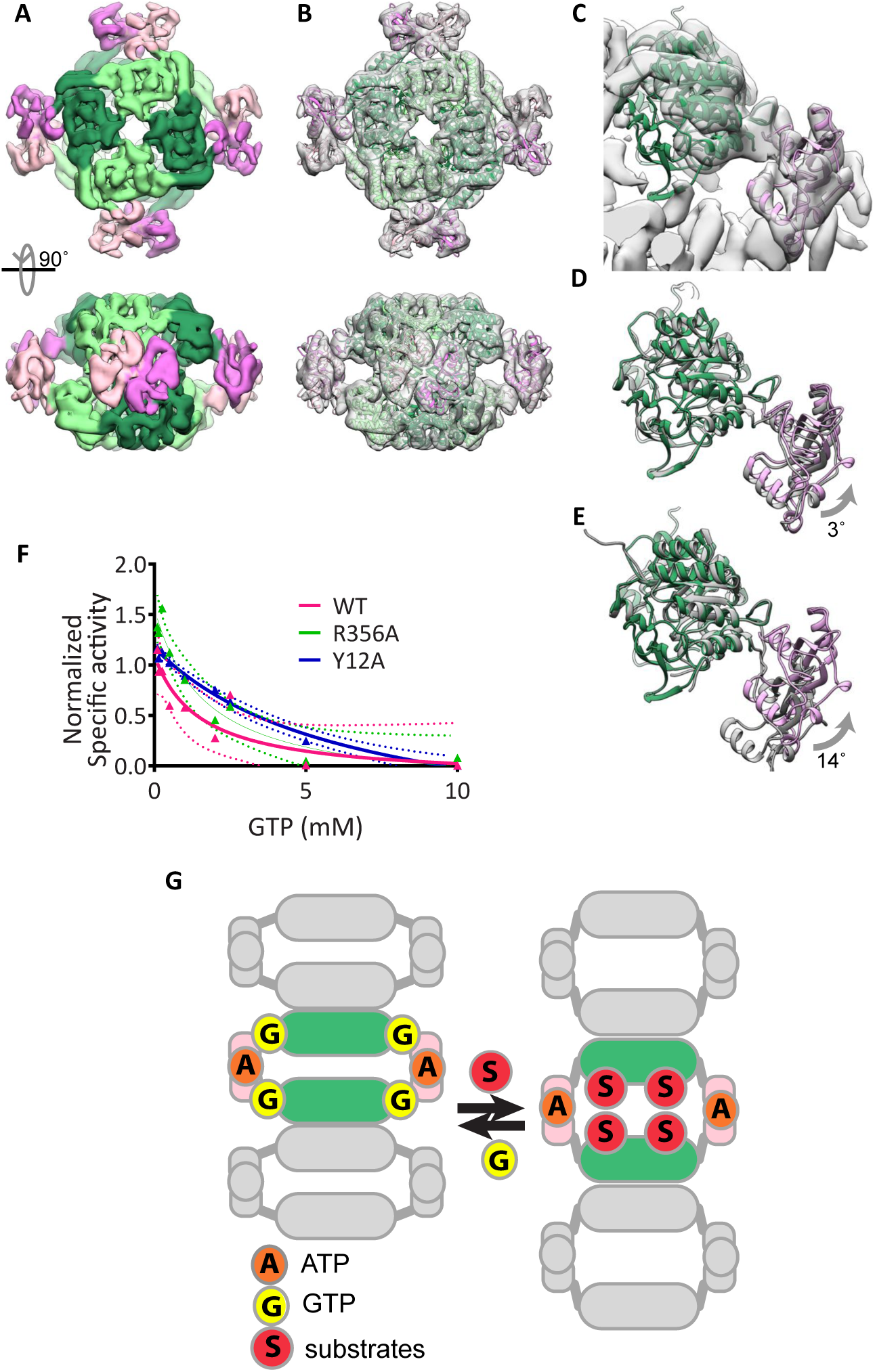
Human IMPDH adopts the collapsed, inhibited conformation when bound to GTP. (A) Cryo-EM reconstruction of human YI2A IMPDH2 non-polymerizing mutant in the presence of 0.1 mM ATP and 0.1 mM GTP, at 8.7 Å resolution. Both catalytic (greens) and Bateman (pinks) domains are well resolved. (B) The atomic model of IMPDH2-ATP-GTP, generated by fitting the catalytic domain and Bateman domains into the cryo-EM structure as two separate rigid bodies. (C) Close-up view of a single IMPDH monomer fit in the cryo-EM structure. (D) A monomer of the A. gossypii IMPDH-GDP crystal structure (gray) (PDB ID 4Z87) aligned to the human IMPDH2-ATP-GTP model (color) on the catalytic domain (green). A rotation of ~3° relates the human Bateman domain (pink) to the A. gossypii structure. (E) The same alignment as (D), but to the P. aueriginosa IMPDH-ATP monomer, in which the Bateman domains are related by a 14° rotation. (F) Dose-dependent inhibition of wild type and mutant IMPDH2 activity by GTP. IMPDH2 was incubated with I mM ATP and 3 mM NAD+ for 10 minutes at room temperature to promote polymerization and then the indicated concentration of GTP was added for an additional 10 minutes prior to reaction initialization by addition of 5 mM IMP. Mean values are plotted and were fit to a three-parameter dose-response curve. Each data point represents 3-6 replicate reactions. 95% confidence intervals are plotted as dotted lines of the corresponding color. (G) Model: human IMPDH exists in a conformational equilibrium between an expanded active conformation and a collapsed inactive conformation that can be shifted by binding to substrates or GTP. Unlike other metabolic filaments, the conformational equilibrium of IMPDH is unaffected by polymerization because the filament form can accommodate both active and inactive conformations.

Since both polymerized and octomeric IMPDH2 can adopt the expanded and collapsed conformations, we hypothesized that polymerization might stabilize the expanded conformation, reducing negative feedback inhibition and allowing higher steady state GTP accumulation in certain biological contexts. To test this, we measured the effect of GTP on the activity of wild type and mutant IMPDH2 under polymerization conditions. Contrary to our hypothesis, we observed comparable low millimolar GTP-mediated inhibition of all three forms (Figure 5F), although we cannot rule out subtle differences in the kinetics or cooperativity of inhibition in the two contexts.

The ability of IMPDH2 filaments to accommodate multiple functional states of the enzyme in different conformations is unexpected. Recent structures of the metabolic filament CTP synthase have shown that, while different ligands influence the conformation of the enzyme, a single conformation is stabilized in the filament, creating a direct link between assembly and enzyme activity (Barry et al. 2014, Lynch et al. 2017). Thus, the ability of IMPDH filaments to accommodate different functionally important conformational states while remaining assembled in the filament is unusual. The conformational change, driven by the balance of substrates and the downstream product GTP, may drive previously unappreciated conformational transitions in macromolecular IMPDH assemblies in vivo.

## MATERIALS & METHODS

### Plasmids and mutagenesis

For expression in mammalian cells, human IMPDH2 was cloned into pcDNA3.1/myc-HIS B. The resulting IMPDH2 protein retains the native N-terminus and has an appended C-terminal myc and 6-histidine tag. Expression is driven by the CMV promoter. Mutagenesis was conducted using QuickChange site-directed mutagenesis (Agilent) according to manufacturer’s instructions. Mutagenesis was conducted with the following forward primers and their corresponding reverse complements:

R356A 5’-CAGAGTATGCACGGGCCTTTGGTGTTCCG-3’

Y12A 5’-GATTAGTGGGGGCACGTCCGCAGTGCCAGACGACGGACTC-3’

S275L 5’-GGATGTAGTGGTTTTGGACTTATCCCAGGGAAATTCCATCTTC-3’

For recombinant IMPDH2 expression in *E. coli*, wild type and mutant IMPDH2 was cloned in pSMT3-Kan (Mossessova and Lima, 2000). This vector appends an N-terminal SMT3/SUMO tag, which can be cleaved by ULP1 protease following purification. As a cautionary note, we found that some N-terminal epitope tags disrupt polymerization of wild type IMPDH2. Consequently, our studies were conducted with IMPDH2 generated with this cleavable tag, which leaves only 5 residual, non-native amino acids. The coding region of all mutated constructs were fully sequenced.

### Cell culture and transfection

HEK293 cells (kindly provided by Dr. Marc Kirschner) were cultured in DMEM (Cellgro) supplemented with 10% FBS (HyClone), 2 mM L-glutamine (Gibco), 1% penicillin and 1% streptomycin. Cells were transfected using Lipofectamine 2000 in Opti-MEM Reduced Serum Medium (ThermoFisher Scientific) as per manufacturer’s instructions.

### Immunoblotting

SDS-PAGE gels were transferred by semi-dry transfer apparatus (Amersham) to nitrocellulose membranes. Membranes were blocked using 5% milk in tris-buffered saline. Anti-myc (Santa Cruz), anti-IMPDH (Abcam) and anti-GAPDH antibodies (Santa Cruz) were used and detected with peroxidase-coupled secondary antibodies and enhanced chemi-luminescence detection (Amersham).

### Immunofluoresence

Cells were gorwn on coverslips and fixed for 20 minutes at room temperature with 4% formaldehyde followed by permeabilitation in 0.5% Triton X-100 for 10 minutes. Blocking and antibody dilution was in phosphate-buffered saline with 0.1% Triton X-100, 2% bovine serum albumin and 0.1% sodium azide. Anti-myc antibodies (Santa Cruz) were used and DAPI was used to counterstain nuclei. Images were collected using MetaVue software (Molecular Devices) controlling a CoolSnap ES camera (Photometrics) on a Nikon TE2000-U inverted microscope using a 60x Plan Apo 1.40 NA Nikon objective.

### Recombinant IMPDH expression and purification

pSMT3-Kan IMPDH2 plasmids were transformed into BL21 (DE3) competent E. coli cells using standard procedures. Transformants were inoculated into 25 ml overnight pre-cultures, which were then used to inoculate 1 L cultures of Luria Broth grown at 37 °C until they reached an OD_600_ of 0.8. Flasks were then cooled on ice for ~5 minutes and then induced with 1 mM IPTG (GoldBio) for 4 hours at 30 °C. Cells were then pelleted for 5 min at 8,000 x g in a Beckmann JA-10 rotor. Cell pellets were stored at -80 °C.

Thawed cell pellets were re-suspended with stirring in 20 ml (per liter of original culture volume) of lysis buffer (50 mM KPO_4_, 300 mM KCl, 20 mM imidazole, 0.8 M urea, pH 8.0) containing Benzonase nuclease (Sigma). Cells were homogenized using the Emulsiflex-C3 (Avestin) with 3 cycles at 10-15 thousand psi. Lysates were then cleared at 18,000 rpm for 20 min at 4 °C in a Beckman Coulter JA 25.5 rotor. SUMO-tagged IMPDH was recovered from the lysates using 1 ml (per liter of original culture volume) of nickel-NTA agarose (Qiagen) and incubation for 2 hours at 4 °C. Beads were washed three times with 10 bead-volumes each of lysis buffer before elution with 50 mM KPO4, 300 mM KCl, 500 mM imidazole, pH 8.0). Peak elution fractions were pooled and incubated with ULP1 protease (purified as described in (Mossessova and Lima, 2000)) (1 mg ULP1 per 100 mg IMPDH2) overnight at 4 °C. 1 mM dithiothreitol (DTT) and 0.8 M urea were added and the protein was concentrated to using a 30,000 molecular weight cutoff Amicon filter and loaded onto either a Superdex 200 or a Superose 6 gel filtration column pre-equilibrated in gel filtration buffer (50 mM Tris, 100 mM KCl, 1 mM DTT, pH 7.4). Peak fractions from the included volume were pooled, aliquotted, flash frozen in liquid nitrogen and stored at -80 °C.

### IMPDH activity assay

NADH production was measured in real time in a 100 μl cuvette (1 cm path length) in a Cary Eclipse fluorometer (excitation: 340 nm, emission: 440 nm) equipped with a temperature controlled multi-cuvette holder maintained at 37 °C. Fluorescence was converted to μmoles NADH produced by comparison with an NADH standard curve. In the standard reaction, 50 mM Tris pH 7.4, 100 mM KCl, 1 mM DTT, 5 mM NAD^+^ and 3 mM IMP. Reactions were initiated by the addition of 13.8 μg of IMPDH2 per 100 μl reaction. Preliminary studies showed these substrate concentrations to be saturating. IMPDH concentration and buffer conditions were chosen to precisely mimic those used for negative stain electron microscopy analysis below. In reactions containing ATP, IMPDH2 was pre-incubated for 10 min at room temperature with the indicated concentration of ATP prior to reaction initiation with substrate. In substrate titration experiments, the constant substrate was included in the pre-incubation and the reaction was initiated by the addition of the variable substrate. In GTP-containing reactions, IMPDH2 was first pre-incubated with 1 mM ATP and 3 mM IMP to induce assembly, then GTP was added to the indicated concentration for an additional 10 minutes before the reaction was initiated by the addition of NAD^+^. Specific activity was calculated by linear interpolation of the reaction slope within the initial 3 min (~ 9 measurements per minute).

### [^13^C_2_, ^15^N]-glycine incorporation into AMP and GMP

HEK293 cells were maintained in DMEM (Cellgro) containing 10% FBS (HyClone) and 2 mM L-glutamine (Gibco). 24 hours prior to the glycine addition, two 10-cm dishes of 40% confluent HEK293 cells were transfected with either the empty vector pcDNA 3.1/myc-His B or S275L IMPDH2 or Y12A IMPDH2 constructs. The next day, cells were washed once in Dulbecco’s PBS (Gibco) and the media of the cells was replaced with DMEM containing 10% dialyzed FBS, 2 mM L-glutamine, and 1.2 mM [^13^C_2_, ^15^N]-glycine (Cambridge Isotope Laboratories). After 6 hours, cells were washed once with Dulbecco’s PBS, released from plates with 0.25% trypsin-EDTA, harvested, washed twice more with Dulbecco’s PBS, and counted. Cells from paired tissue culture dishes were pooled, resulting in 1-10x10^6^ cells per sample.

Samples were then processed and analyzed as described previously (Laourdakis *et al.*, 2014). Briefly, 70 μL of a solution containing 0.5 M perchloric acid and 200 μM of the internal standard [^13^C_9_, ^15^N_3_]-CTP was added to each cell pellet. The cells were vortexed for 10 seconds and incubated on ice for 20 minutes. Lysates were then neutralized with 7 μL of 5 M potassium hydroxide, vortexed for 10 seconds, and incubated on ice for an additional 20 minutes. Cell debris was harvested by centrifugation at 11,000 x g for 10 minutes at 4 °C. Supernatants were further clarified using Amicon Ultra 0.5 mL centrifugal filters (3 kD MWCO) as per manufacturer’s instructions.

Collected extracts were analyzed by an ultra performance liquid chromatography (UPLC, Waters Acquity I-class) coupled to a triple quadrupole mass spectrometer (Waters Xevo TQ-S micro). A Waters Acquity UPLC HSS T3 column (1.8 mm, 2.1 mm x 50 mm) and a column in-line filter were used. Quantitative analysis of nucleotides was performed using the multiple reaction monitoring mode, and the peaks of nucleotides were analyzed using MassLynx V4.1. In addition, calibration standards were freshly prepared for every sample set analysis, and linear calibration curves were established. All reported data represent the mean and standard error of three independent biological replicates. *P* values were calculated using two-tailed, unpaired Student’s *t* tests in Graphpad Prism.

For each replicate, samples for immunofluorescence and immunoblotting were prepared in parallel to samples for nucleotide extraction. Cells for immunofluorescence were stained as described above except that DRAQ5 (Biolegend) was used as the nuclear stain. Images were obtained using a Leica DM 5000 microscope coupled to a Leica TCS SP5 confocal laser scanner. For immuno blot analysis, cells were lysed in RIPA buffer supplemented with Complete protease inhibitor cocktail (Roche). Proteins were separated by SDS-PAGE and transferred to nitrocellulose. Blots were stained with anti-IMPDH2 antibody EPR8365(B) (Abcam).

### Negatively stained electron microscopy and particle averaging/reconstruction

Aliquots of purified protein, including WT IMPDH and the mutants Y12A, R356A, and S275L, were diluted to 2.5 μM in assembly buffer (50 mM Tris, 100 mM KCl, 1 mM DTT, pH 7.4) and incubated for 15 minutes at room temperature in the presence of various concentrations of ATP, GTP, IMP, and NAD^+^, as outlined in figures 1D and 4C. Following incubation, samples were applied to glow-discharged continuous carbon film EM grids and negatively stained using 1% uranyl formate. Grids were imaged by transmission electron microscopy using an FEI Tecnai G2 Spirit operating at 120 kV and a Gatan Ultrascan 4000 CCD via the Leginon software package (Suloway *et al.*, 2005). Image processing, including CTF estimation, particle picking, and two-dimensional reference-free classification, was performed using the Appion and RELION software packages (Lander *et al.*, 2009; Scheres, 2012). The image data collected from all ATP/GTP/IMP/NAD^+^ combinations were combined and processed *en masse* (77,525 helical repeats in total). Following 2D classification, the resulting class averages were manually grouped according to the observed rise into one of three categories: “compressed” (rise ~94 Angstrom), “extended” (rise ~114 Angstrom), or “undetermined” (filament ends, distorted and bent octamers, and poorly aligned particles). The proportion of helical repeats from each experimental dataset belonging to these three categories was then back-calculated.

For three-dimensional reconstructions of IMPDH filaments, negative stain images were acquired at a pixel size of 2.07 Å and individual filaments were identified manually in Appion (Lander *et al.*, 2009). Reconstructions were performed by iterative helical real space refinement (Egelman, 2010) in SPIDER(Shaikh *et al.*, 2008), imposing D4 symmetry on the helical asymmetric unit throughout. The structures of the collapsed and expanded IMPDH filament have been deposited in the EMDB (Lawson *et al.*, 2011) (EMD-8691, expanded, and EMD-8690, collapsed).

### Electron cryo-microscopy and particle averaging/reconstruction

Purified Y12A protein was diluted in assembly buffer to 4 uM, and incubated at room temperature in the presence of 0.1 mM ATP and 0.1 mM GTP for 15 minutes. Following incubation, the sample was applied to a glow-discharged C-Flat holey carbon EM grid (Protochips), and blotted/plunged into liquid ethane using a Vitrobot plunging apparatus (FEI). Imaging was performed using an FEI Tecnai G2 F20 operating at 300 kV and a Gatan K-2 direct electron detector via the Leginon software package (CITE Suloway 2005, JSB). Image processing, including CTF estimation, particle picking, two-dimensional reference-free classification, three-dimensional classification and 3D refinement, was performed using the Appion and RELION software packages (CITE Lander 2009, JSB & Scheres 2012 JSB). The map resolution, as determined by the relion_postprocess script using the Fourier shell correlation (FSC) = 0.143 criterion was 8.7 angstroms. The final electron density map was further sharpened using a b-factor of -987. The structure has been deposited with the EMDB (EMD-8692).

### Statistics

Statistical tests used are indicated in the figure legends and *P* values < 0.05 were considered significant.

## ACKNOWLEDGEMENTS

This work was supported by National Institutes of Health R01 grants GM083025 to J.R.P., GM118396 to J.M.K., and GM102503 to A.A. K.C.D was supported, in part, by T32 CA009035. We thank Alana O’Reilly for the use of her confocal microscope and insightful discussions and Liz Hedstrom for IMPDH expression constructs.

---

**Supplemental Figure 1.**
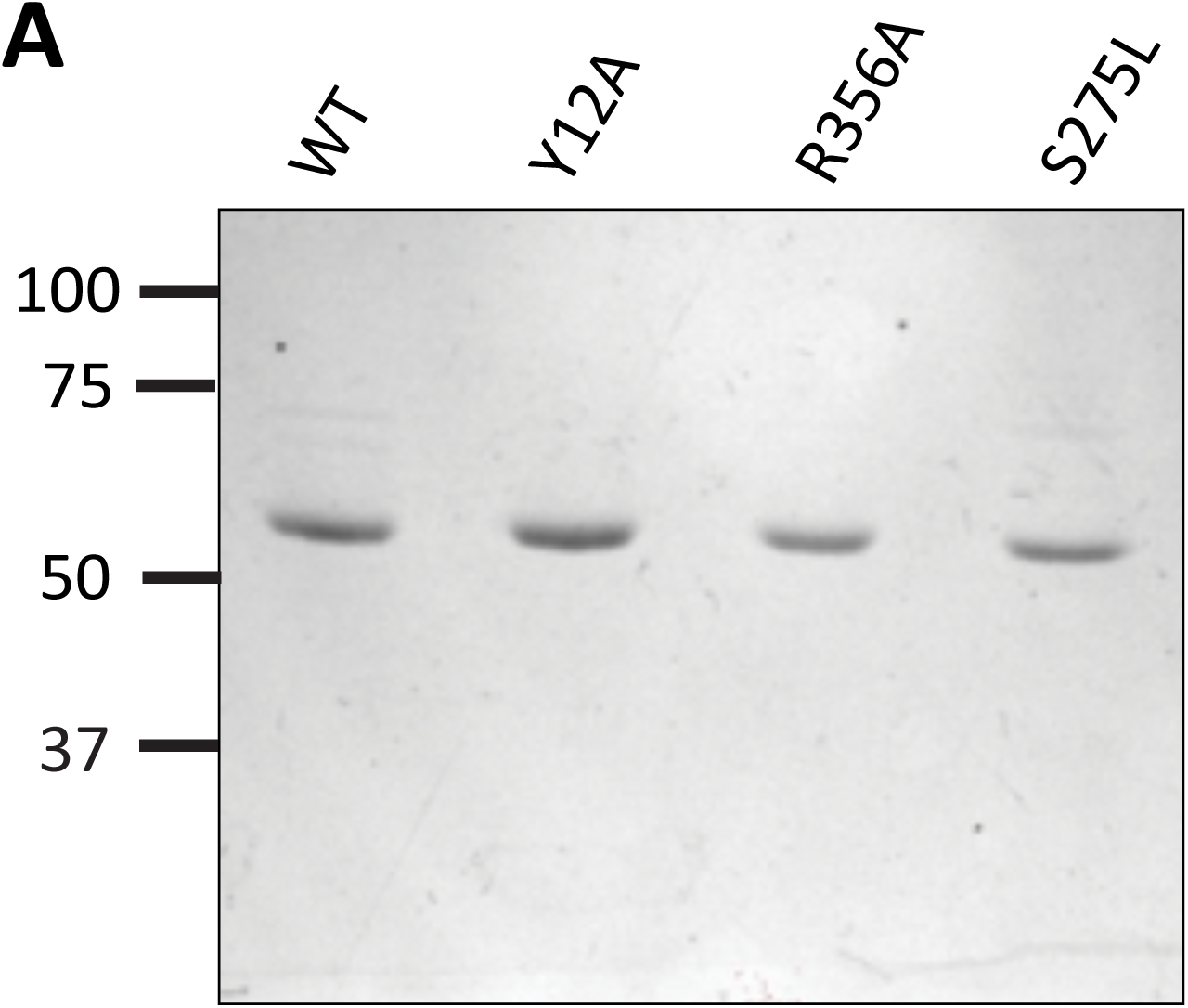
Coomassie-stained gel of purified IMPDH2 wild type and mutant proteins.

**Supplemental Table 1:** Michaelis-Menten parameters derived from substrate titrations in Figure 3.

|  | **IMP** |  |  |  |
| --- | --- | --- | --- | --- |
|  | **Y12A** | **R356A** | **WT** | **WT no ATP** |
| **normalized Vmax**<br>($\mu\text{mol}/\text{min}/\text{mg}$) | 0.9963 | 0.9961 | 1.04 | 0.9996 |
| **K<sub>m</sub> ($\mu\text{M}$)** | 0.01535 | 0.02153 | 0.01295 | 0.01749 |
| **95% CI (profile likelihood)** |  |  |  |  |
| **Vmax** | 0.9362 to 1.059 | 0.8906 to 1.111 | 0.9251 to 1.161 | 0.839 to 1.174 |
| **K<sub>m</sub> ($\mu\text{M}$)** | 0.01109 to 0.02087 | 0.01123 to 0.03932 | 0.006642 to 0.02349 | 0.006703 to 0.03968 |
|  | **NAD<sup>+</sup>** |  |  |  |
|  | **Y12A** | **R356A** | **WT** | **WT no ATP** |
| **normalized Vmax**<br>($\mu\text{mol}/\text{min}/\text{mg}$) | 1.001 | 1.008 | 1 | 1.079 |
| **K<sub>m</sub> ($\mu\text{M}$)** | 0.01107 | 0.005968 | 0.007649 | 0.0007609 |
| **95% CI (profile likelihood)** |  |  |  |  |
| **Vmax** | 0.8868 to 1.121 | 0.8755 to 1.149 | 0.8025 to 1.218 | 0.9778 to 1.183 |
| **K<sub>m</sub> ($\mu\text{M}$)** | 0.005614 to 0.01988 | 0.001903 to 0.01299 | 0.002003 to 0.01879 | 0.0002202 to 0.002949 |

